# The recovery of plant community composition following passive restoration across spatial scales

**DOI:** 10.1101/2022.04.14.488351

**Authors:** Emma Ladouceur, Forest Isbell, Adam T. Clark, W. Stanley Harpole, Peter B. Reich, G. David Tilman, Jonathan M. Chase

## Abstract

1. Human impacts have led to dramatic biodiversity change which can be highly scale-dependent across space and time. A primary means to manage these changes is via passive (here, the removal of disturbance) or active (management interventions) ecological restoration. The recovery of biodiversity, following the removal of disturbance is often incomplete relative to some kind of reference target. The magnitude of recovery of ecological systems following disturbance depend on the landscape matrix, as well as the temporal and spatial scales at which biodiversity is measured.
2. We measured the recovery of biodiversity and species composition over 27 years in 17 temperate grasslands abandoned after agriculture at different points in time, collectively forming a chronosequence since abandonment from one to eighty years. We compare these abandoned sites with known agricultural land-use histories to never-disturbed sites as relative benchmarks. We specifically measured aspects of diversity at the local plot-scale (α-scale, 0.5m^2^) and site-scale (γ-scale, 10m^2^), as well as the within-site heterogeneity (β-diversity) and among-site variation in species composition (turnover and nestedness).
3. At our α-scale, sites recovering after agricultural abandonment only had 70% of the plant species richness (and ∼30% of the evenness), compared to never-ploughed sites. Within-site β-diversity recovered following agricultural abandonment to around 90% after 80 years. This effect, however, was not enough to lead to recovery at our γ-scale. Richness in recovering sites was ∼65% of that in remnant never-ploughed sites. The presence of species characteristic of the never disturbed sites increased in the recovering sites through time. Forb and legume cover declines in years since abandonment, relative to graminoid cover across sites.
4. ***Synthesis.*** We found that, during the 80 years after agricultural abandonment, old-fields did not recover to the level of biodiversity in remnant never-ploughed sites at any scale. β-diversity recovered more than α-scale or γ-scales. Plant species composition recovered, but not completely, over time, and some species groups increased their cover more than others. Patterns of ecological recovery in degraded ecosystems across space and long time-scales can inform effective, targeted active restoration interventions and perhaps, lead to better outcomes.

## Introduction

The Anthropocene is characterised by dramatic impacts of people on the biosphere, via a number of direct and indirect processes (e.g., land use, climate change), often leading to altered numbers and types of species (i.e., biodiversity) in those impacted ecosystems (Díaz et al., 2019; Newbold et al., 2015). While a primary means to manage these changes is to reduce the extent and intensity of negative drivers of biodiversity change (e.g., reduced destruction or degradation of natural ecosystems), an increasingly important way to recover losses of biodiversity and the ecosystem services it provides is via a cessation or reduction of the impacts and restoration of those ecosystems (Jones et al., 2018). The United Nations has recently announced 2021-2030 as the Decade on Ecosystem Restoration, with the goal of restoring 350 million hectares of degraded land to achieve higher biodiversity and ecosystem functions (UNEA, 2019).

Ecological restoration can take many forms. The Society for Ecological Restoration (SER) recognises a ‘restorative continuum’ of interventions that can help ecosystems recover to context-dependent benchmarks after disturbance (Gann et al., 2019). This can range from passive restoration, or natural recovery, which is the cessation of major disturbance (e.g., deforestation, agriculture) (Atkinson & Bonser, 2020; Chazdon et al., 2021) or reinstatement of disturbance and management regimes (e.g., fire, grazing, mowing). Active, or assisted and reconstructive restoration, includes the addition of interventions which might manipulate abiotic and biotic factors including the reintroduction of desired biota (Atkinson & Bonser, 2020). Through successional processes or the assistance of such processes, such as recolonizations and extinctions, ecosystems can then recover on a trajectory towards a desirable functioning state (Shackelford, Dudney, et al., 2021; Temperton et al., 2004). However, this recovery is typically incomplete (Jones et al., 2018; Moreno-Mateos et al., 2017; Rey Benayas et al., 2009). In addition, communities in restored/recovered ecosystems are often composed of more generalist and alien species when compared to reference sites (Kaul & Wilsey, 2021). The composition of recovering sites at different time points can be tied to species life-history characteristics (Zirbel & Brudvig, 2020), can have interactive inhibitory or facilitative effects for other species to recolonise (Young et al., 2017), and can be influenced by the surrounding landscape, history, and management (Funk, 2021; Grman et al., 2015; Guiden et al., 2021).

Despite frequent studies on how biodiversity responds to anthropogenic impact and recovery (Murphy & Romanuk, 2014; Newbold et al., 2015), less attention has been paid to how inference of restoration on biodiversity depends on the ecological scale in which diversity is measured and observed (Catano et al., 2021; Martin et al., 2005). Nevertheless, most measures of biodiversity inherently depend on the spatial scale on which samples are taken (i.e., a 1m^2^ quadrat compared to an entire site), and on the temporal scales which are measured (i.e., a year or a decade) (Matthews et al., 2021; Rosenzweig, 1995). As a result, scale can critically influence the magnitude in which biodiversity changes are quantified, even when sample effort is standardised (Chase et al., 2018; Chase & Knight, 2013; Field et al., 2009; Hill & Hamer, 2004; Sax & Gaines, 2003).

While the scale-dependence of biodiversity responses to anthropogenic activities are well known, the direction of scale dependence is less clear. Scale-dependent biodiversity responses to anthropogenic activities are most often studied in the context of changes to β-diversity, or the site differences in species composition (Chase et al., 2019; Socolar et al., 2016). Often, anthropogenic activities are thought to create a homogenising effect, reducing β-diversity (Gossner et al., 2016; Hautier et al., 2018; Martin et al., 2005). When β diversity is reduced by an anthropogenic driver, this can lead to cases where small to moderate effects of a driver at smaller scales (i.e., α-diversity) can become exacerbated at larger spatial scales. For example, (W. Li et al., 2021) found that Mongolian semiarid grassland communities that were impacted by grazing and mowing had fewer species in each locality (α-diversity) as disturbance intensity increased. However, because more narrowly distributed species were more strongly influenced by disturbance intensities than more widespread species, β-diversity also decreased, and the effect at the larger (γ-diversity) spatial scale was greater.

On the other hand, anthropogenic activities can lead to communities becoming more different between sites (higher β-diversity), which can lead to cases where relatively larger effects of a driver occur at smaller spatial scales (i.e., α-diversity). These negative effects weaken as scale increases (i.e., γ-diversity). For example, (Uchida et al., 2018) found that land abandonment in Japanese semi-natural grasslands led to a reduction in small-scale species richness when compared to intensive agriculture and traditional management practices, but this negative effect sometimes dissipated as scale increased. Likewise, semi-natural grassland communities in the Slovak Republic had lower α-diversity, but an increase in β-diversity and γ-diversity in landscapes with a higher proportion of non-natural habitats (Janišová et al., 2014). There are many underlying factors that can influence the direction and magnitude of scale-dependence resulting from anthropogenic drivers, with numerous examples supporting each (Chase et al. 2018, 2019).

Spatial scale can influence our understanding of how biodiversity recovers following cessation of major disturbance. For example, small-scale (α) diversity usually does not fully recover to pre-disturbance levels even under active restoration (Isbell et al., 2019; Moreno-Mateos et al., 2017; Rey Benayas et al., 2009). What is less clear, however, is how β and γ-diversity respond during recovery. If β-diversity is not influenced by the removal of disturbance and does not increase through time, the incomplete recovery of diversity following restoration would be equivalent at both α (within-plot) and γ (site) scales. If β-diversity is reduced by removal of disturbance (i.e., via homogenization) and does not recover during restoration, the incomplete recovery of α diversity would be exacerbated at larger (γ) scales. Finally, if β-diversity increases following removal of the disturbance, incomplete recovery of α diversity could be accompanied by a more complete recovery of diversity at larger spatial scales. To date, few studies have examined the influence of recovery and restoration on the scale-dependence of biodiversity and β-diversity in particular, and those that have measured β-diversity do so in a number of different, often non-comparable, ways. There is some evidence that β-diversity and γ-diversity recover even less than α-diversity (Martin et al., 2005; Passy & Blanchet, 2007; Polley et al., 2005; Wilsey et al., 2005). In a meta-analysis of grassland studies, however, (Catano et al., 2017) showed that the effects of disturbance can lead to homogenization (lower β-diversity) or differentiation (higher β-diversity) depending on the effects of disturbance on stochastic factors and dispersal rates. Furthermore, β-diversity can be enhanced in restoration, for example, when restoration actively targets β-diversity via larger species pools (Grman & Brudvig, 2014).

While there is evidence of deficits of α, β and γ-diversity in passively recovering and actively restored ecosystems, and in grassland systems in particular (Martin et al., 2005; Polley et al., 2005; Sluis, 2002), it remains unclear how long these potential deficits manifest on the landscape. Grasslands are one of the most endangered and least protected biomes globally (Hoekstra et al., 2004), and are experiencing extreme levels of land use change locally and regionally (Carbutt et al., 2017; Roch & Jaeger, 2014). There are few old growth and continuous tracts of grassland left (Nerlekar & Veldman, 2020; Scholtz & Twidwell, 2022). Here, we take advantage of long-term surveys of vegetation in remnant savannah prairies and recovering grasslands at the Cedar Creek Ecosystem Science Reserve in Minnesota (USA). Isbell et al. (2019) used data from remnant prairies and a 37-year survey of old-fields with different amounts of time since agricultural abandonment (ranging from 1 to 91 years) to examine how α-scale (within-plot) species richness, species diversity, evenness and productivity recovered (measured in 0.3 m^2^ plots). They found that even after more than 91 years since abandonment of agriculture, species richness only recovered to 75% of its value in the reference site that was never-ploughed. However, because species richness is a scale-dependent metric, it is unclear how larger scale diversity recovered. Using the same system, and some, but not all of the same sampling as Isbell et al. (2019), and taking different spatial scales explicitly into account, we asked the following: (1) how do larger scale measures of diversity (i.e., β and γ-diversity) vary through time across the chronosequence following agricultural abandonment and how do they compare to the smaller scale (α) measures? (2) How do measures of diversity other than species richness, such as those that incorporate evenness, respond at the α, β and γ-scales? (3) How has species composition, as a different component of recovery compared to measures of richness and diversity, responded through time? (4) How has the cover of species with different growth forms and life histories responded through time?

## Methods

### Study system

Cedar Creek Ecosystem Science Reserve, hereafter referred to simply as Cedar Creek, is a 2,200-hectare long-term ecological science reserve (LTER) run by the University of Minnesota (USA) in cooperation with the Minnesota Academy of Science located 50 km north of Minneapolis. Cedar Creek lies on a glacial outwash sand plain, between deciduous forest to the east and prairie to the west, forming a mosaic of oak savanna, prairie, upland deciduous forest, lowland marshes and swamps (Inouye et al., 1987). Soils are largely outwash sediments of fine and medium sands, poor in nitrogen, which was further depleted by agricultural practices in old field sites (Inouye et al., 1987).

Agricultural land use in this area began after 1900, but aerial photography suggests some areas were never cleared (MHAPO, 2015; Pierce, 1954). As a result, there are now a series of agricultural sites (old-fields) abandoned at different times during the last century under passive recovery, as well as never-ploughed remnant prairies and savannas scattered across the reserve (Fig. S1, Table S1). Secondary succession in the abandoned old-fields is significantly limited by nitrogen (Tilman, 1987), and dispersal limitation (Tilman, 1994). While fire does play a key role in maintaining prairies and savannas, succession does not consistently lead to afforestation in the absence of fire (Clark et al., 2019). The natural history of Cedar Creek is described in more detail in (Inouye et al., 1987).

### Study design and sampling

We analysed vegetation from several sites that were part of long-term research at the Cedar Creek Ecosystem Science Reserve (Fig. S1). Specifically, we used data from otherwise comparable sites that could be categorised into two states (see Table S1 for details): (1) Never-ploughed sites, which included 18 upland oak savannas (plots of “Experiment 133”); (2) recovery sites that included 17 old-fields which were ploughed and used for agriculture, but were abandoned so that natural succession and recovery of the vegetation could be followed (“Experiment 014”). Old-fields were abandoned between 1927 and 2015 (Clark et al., 2019). Each field was measured approximately 6 times, with ∼5-6 year measurement intervals from 1983-2010 (27 years). At the start of the surveys, old-fields ranged from 1 year since agricultural abandonment to 48 years (Table S1). All sampled sites were located on well-drained upland sands (Inouye et al., 1987).

For surveys in both experiments, plants were estimated using percent cover classes (“Experiment 133”) and species-level percent cover (“Experiment 14”) in 0.5 m^2^ plots (1m x 0.5m). Cover classes are based on a modified Domin scale (1 = 1%, 2 = 2-5%, 3 = 6-25%, 4 = 26-50%, 5 = 51-75%, and 6 = 76-100%). Cover in both studies could exceed 100%. In Experiment 133 (never-ploughed fields), four parallel 50 m long transects were laid out within fire management block units within each field, 25 m apart, and 6 plots were placed every 10m along each transect, for a total of 24 plots in fields in most years. In Experiment 14 (old fields), four permanent parallel 40m long transects were laid in each field, 25 m apart, and 25 plots were placed every 1.5 m along each transect, totalling 100 plots per field.

We did not include plots from the never-ploughed sites that had also never been burned. This is because fire is a natural disturbance that maintains these systems, and woody encroachment ensues when there is human-induced fire suppression (Clark et al., 2019). Within the old-fields, we kept all plots surveyed in years before the first year of burning, or those that have not been burned to represent site recovery after abandonment before fire. We also did not include sites that contained many trees (Clark et al., 2019). Because sample effort was not equal between all of the sites in some years (e.g.,1999, 2011), we selected sites and years that had a minimum of 24 samples, and used 20 randomly selected survey plots from each site (site E14 was sub-sampled to match the minimum number of samples in site E133). We took the midpoint of each cover class (“Experiment 133”) to quantify percent cover of species, so that the summed cover of all species could exceed 100%. Species cover relative to each plot’s summed cover was quantified as a proportion.

### Calculating Within-Site Metrics of Diversity

We examined how biodiversity recovered across scales by calculating and comparing multiple metrics of diversity at multiple scales between the never-ploughed and recovery treatments. We estimated diversity at two spatial scales: (i) the α-scale, which was the diversity in a given 0.5m^2^ plot in a given treatment and year, and (ii) the γ-scale, which here we define as the total diversity in 20 0.5m^2^ of combined plots within a site and year. Note, here we simply use α and γ-diversity to denote smaller and larger scales, and make no assumptions whether these scales correlate with any local or regional coexistence mechanisms. Finally, from these α and γ estimates, we calculate (iii) Whittaker’s multiplicative β-diversity (β = γ/α, (Whittaker, 1972) to quantify plot-to-plot variation, or the heterogeneity of plots within sites at each time point. While the sampling approach was not designed to sample the range of variation in the whole site, the amount and the equal number of samples across sites allows an estimation of this variation. Additionally, we extrapolated expected species richness from the γ-scale across 50 samples (Chao et al., 2014).

At each observed spatial scale, we estimated two metrics of diversity: (i) species richness, which was simply the total number of species observed in a given α-plot or γ-site, and (ii) an estimate of diversity that more heavily weights common species, the probability of interspecific encounter (PIE). The PIE is the probability that two species sampled randomly from a community are of a different species (Hulbert, 1971), and higher values represent more even communities. For analyses, we transformed the PIE into an effective number of species (ENS_PIE_), that has the same number of units as species richness using the proportion of each species (Jost, 2006); this is equivalent to Simpson’s inverse diversity index (Simpson, 1949; Williams, 1964). By comparing results of species richness versus ENS_PIE_, we can evaluate whether differences are more strongly influenced by rare species only (in which case species richness results should be different from ENS_PIE_ results), or by both rare and common species (in which case, results from both metrics would be more similar) (Smith & Wilson, 1996). At all scales, we standardized the cover to sum 100%. At the α-scale, we summed cover across all species, and at the γ-scale, we summed across all species and plots. We used this relative proportion to calculate ENS_PIE_ at all scales.

### Species composition

The measures of biodiversity (species richness and ENS_PIE_) explored here across scales (α, β, γ) allow us to compare numbers and types of species from plots within a given site status (i.e., never-ploughed versus ploughed sites). They do not, however, allow us to quantify the difference in species composition between the sites. For example, many highly specialised prairie and savanna plants rarely establish in the early phase of old-field recovery, which instead is dominated by weedy species that are less frequently found in pristine sites (Inouye et al., 1987).

To quantify the difference in species composition between never-ploughed sites and recovering sites, we calculated Jaccard’s dissimilarity index indicating the dissimilarity in species composition between site status (ranging from 0 to 1). We partitioned the difference between the turnover and nestedness components of Jaccard’s index (Baselga, 2009; Baselga & Orme, 2012). These metrics are known to be sensitive to pre-existing differences in alpha-diversity (Vellend et al., 2007). Through the careful design of our approach, we avoid these issues.

We used a checklist approach to identify species present within old-fields and across fields that have never been ploughed. First, we compiled a checklist of all species present across all never-ploughed sites within every time point measured (1984, 1990, 1995, 2000, 2005, 2010) to determine the ‘regional’ species pool for the never-ploughed sites within each year, resulting in one ‘regional checklist’ for every year. This never-ploughed regional checklist represents a temporally accurate relative restoration reference target. Next, we compiled a checklist of all species present within every old-field site and time point measured (1983, 1989, 1994, 1997, 2002, 2006), resulting in a checklist for every old field site and every year. If a species was present across multiple plots within a site, it was only represented once in the never-ploughed or old-field checklist with a presence of 1. We then compared species present in the checklist of every old-field site within each year to the regional checklist across all never-ploughed sites at the closest calendar year and nearest comparable time-point measured (eg. 1983 compared to 1984) as a single pairwise comparison (Marion et al., 2017). This quantifies the site-level compositional change of each old-field since agricultural abandonment relative to the total species present within all never-ploughed sites at the most comparable time-point as a consistent comparison benchmark. If species from the checklist of never-ploughed sites were recolonizing old-fields over time, we expect nestedness to increase in each old-field as years since agricultural abandonment progress. If species colonising old-field sites since abandonment are different from that of never-ploughed sites, then we would expect the turnover component to increase across time.

Lastly, to quantify changes in relative cover after years since agricultural abandonment, we calculated the relative cover of species broad growth form groups (graminoids, forbs, and legumes) and their origin (native, exotic) for every plot measured in every year.

We used the R Environment for Statistics and Computing (R Core Development Team, 2019) for all data preparation, manipulation, quantification of diversity metrics, statistical analysis and graphic visualisation of results. To quantify diversity metrics (e.g., α, β, γ species richness) we used tidyverse (Wickham et al., 2019) and vegan (e.g., α, β, γ ENS_PIE_) packages (Oksanen et al., 2019). For dissimilarity and its partition into turnover and nestedness, we used the beta.part package (Baselga & Orme, 2012; Oksanen et al., 2019). We used the iNEXT package (Chao et al., 2014; Hsieh, T.C. et al., 2020) to interpolate and extrapolate (up to 50 samples) species richness across scales based on average observed samples across each site status and year since agricultural abandonment (Fig. S2, S3).

### Statistical analysis

We quantified field status (old-field vs never-ploughed) effects on biodiversity using hierarchical linear models (Discrete analysis) with site status (i.e., old-field or never-ploughed) as a categorical fixed effect. We modelled site and calendar year as random effects, and allowed random intercepts to vary (Supplementary Information Table S2, Fig. S4-S9 for all model details).

The effect of years of agricultural abandonment on the recovery of old-fields compared to the never-ploughed sites at each scale of diversity were quantified using hierarchical linear models (continuous analysis). Year since agricultural abandonment was modelled as a continuous fixed effect and was log-transformed. We found logarithmic trends to be the most parsimonious for statistical models looking at diversity (e.g, α, β, γ, richness and ENS_PIE_) as a function of years since abandonment, similar to previous work (Isbell et al., 2019). At both spatial scales, we quantified each log-transformed diversity component in old-fields as the percentage compared to the mean of the never-ploughed sites at that same metric and scale.

We allowed random intercepts and slopes to vary for years since agricultural abandonment, and for categorical calendar year, assuming variation between each site, the year it was abandoned and the calendar year sampling started (Supplementary Information for all model details). For the α-scale, we modelled plot nested within transect, nested within site as random effects and for β and γ-scales, site as a random effect. We plotted log-transformed trends on a linear scale, which is why the visualisation of the overall trends have some curvature.

We quantified the year of abandonment effects on the turnover and nestedness components of Jaccard’s dissimilarity using two univariate hierarchical linear models (dissimilarity analysis). We modelled the year since abandonment as a continuous fixed effect and site as a random effect. We allowed random intercepts and slopes to vary for years since agricultural abandonment, for every old field and for calendar year, assuming there was variation in each site and each year of sampling We quantified the effect of years since agricultural abandonment on the relative cover of growth forms (graminoid, forb, legume) and their origin (native and introduced) using a univariate hierarchical linear model. We modelled the year since abandonment as a continuous fixed effect (log-transformed), including growth form and origin and their 3-way-interaction as categorical fixed effects. We log-transformed relative percent cover and allowed random intercepts and slopes to vary for all fixed effects across every site, and year. We plotted log responses on a linear scale.

For Bayesian inference and estimates of uncertainty, we fit models using the Hamiltonian Monte Carlo (HMC) sampler Stan (Carpenter et al., 2017), and coded using the brms package (Bürkner, 2018). We fit all models with four chains, differing iterations, and differing assumed distributions (See Supplementary Information for all model details). We used weakly regularising priors and visual inspection of the HMC chains showed excellent convergence.

## Results

### α-scale and γ-scale species richness

For the discrete analysis (Figs 1 a, b), we found that, old-field plots had on average 59% of species richness (5.65, 95% credible interval: 4.66 to 6.56) found in never-ploughed sites (9.59, CI: 8.70 to 10.51, Fig. 1a) at the α-scale (0.5 m^2^) and approximately 55% fewer species (23.6, 21.2 to 26.1) than never-ploughed sites (42.6, 38.6 to 47.1, Fig. 1b) at the γ-scale (the combination of 20 plots and thus 10m^2^). We found a similar difference between old-fields and never-ploughed sites when we extrapolated species richness estimates based on incidence-based species accumulation to 50 samples (Figs S2, S3).

In the continuous analysis (Figs 1 c, d), richness weakly increased across years since agricultural abandonment at the α-scale, but with high uncertainty (Slope: 0.13, 95% credible interval: -0.17 to 0.53, Fig. 1c). After ∼80 years since abandonment, α-scale species richness in old-fields was about 70% of that found in never-ploughed sites. At the γ-scale, grassland species richness increased across years since agricultural abandonment (Fig. 1d), but again with high uncertainty (Slope: 0.13, CI: -0.11 to 0.40). γ-scale richness in old-fields was about 65% of that found in never-ploughed sites after 80 years.

**Figure 1:**
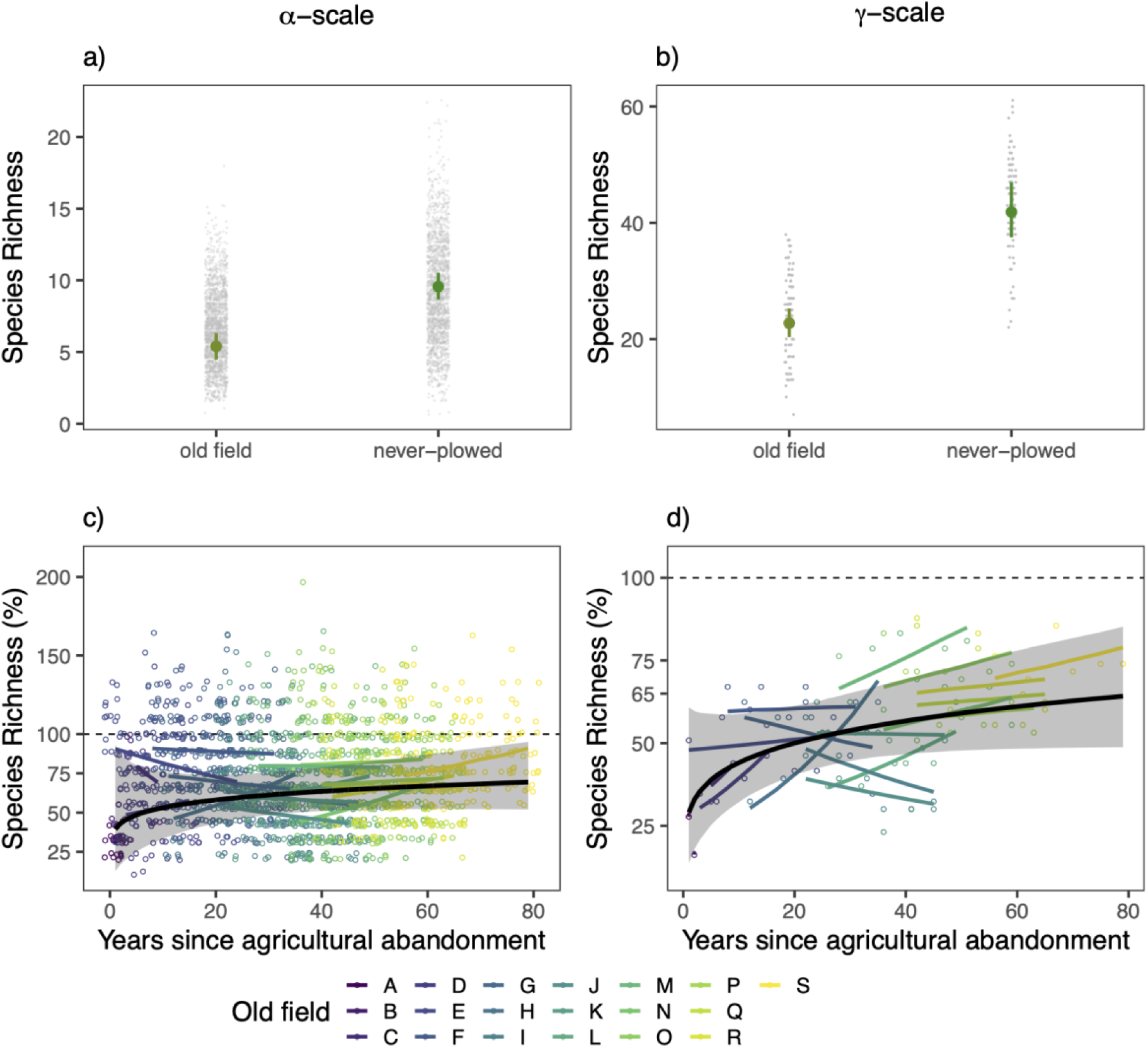
α-scale and γ-scale species richness. a) α-scale and b) γ-scale species richness as a function of site status. Small points show data models were fit to; large points are the conditional effects of site status and the lines show the 95% credible intervals of conditional effects. c) α-scale and d) γ-scale species richness as a function of ‘years since agricultural abandonment’. Black dashed line represents the mean diversity metric of all never-ploughed sites (18 sites). The thick black line represents the average effect of years since agricultural abandonment on species richness across all old-fields. The grey shading around the black line represents the 95% credible interval of that effect estimate. Each colored line represents the average predicted values for each site. Each open point represents an old-field plot (α-scale) or a site (γ-scale) calculated as the percentage of richness, compared to never-ploughed sites. Y-axes vary for clarity.

### α-scale and γ-scale ENS_PIE_

In discrete analysis (top row) at the α-scale, old-field sites overall had approximately 31% of ENS_PIE_ (2.27, 1.66 to 2.89) (relative abundance was less even) found in never-ploughed sites (7.21, 6.62 to 7.82, Fig. 2a). At the γ-scale, old-field sites had approximately 28% (3.53, 2.14 to 4.88) fewer species equivalents than in never-ploughed sites (12.61, 11.27 to 14.02, Fig. 2b).

At the α-scale, ENS_PIE_ does not increase strongly across years since agricultural abandonment (Slope: 0.07, 95% Credible Intervals: -0.14 to 0.37, Fig. 2c). After ∼ 80 years, α-scale plots within old-fields have less than 50% ENS_PIE_ than those that were never disturbed. At the γ-scale, ENS_PIE_ weakly increases across years since agricultural abandonment with high uncertainty (Slope: 0.15, CI: -0.08 to 0.36, Fig. 2d), and had less than 50% ENS_PIE_ than the never-ploughed sites after 80 years.

**Figure 2:**
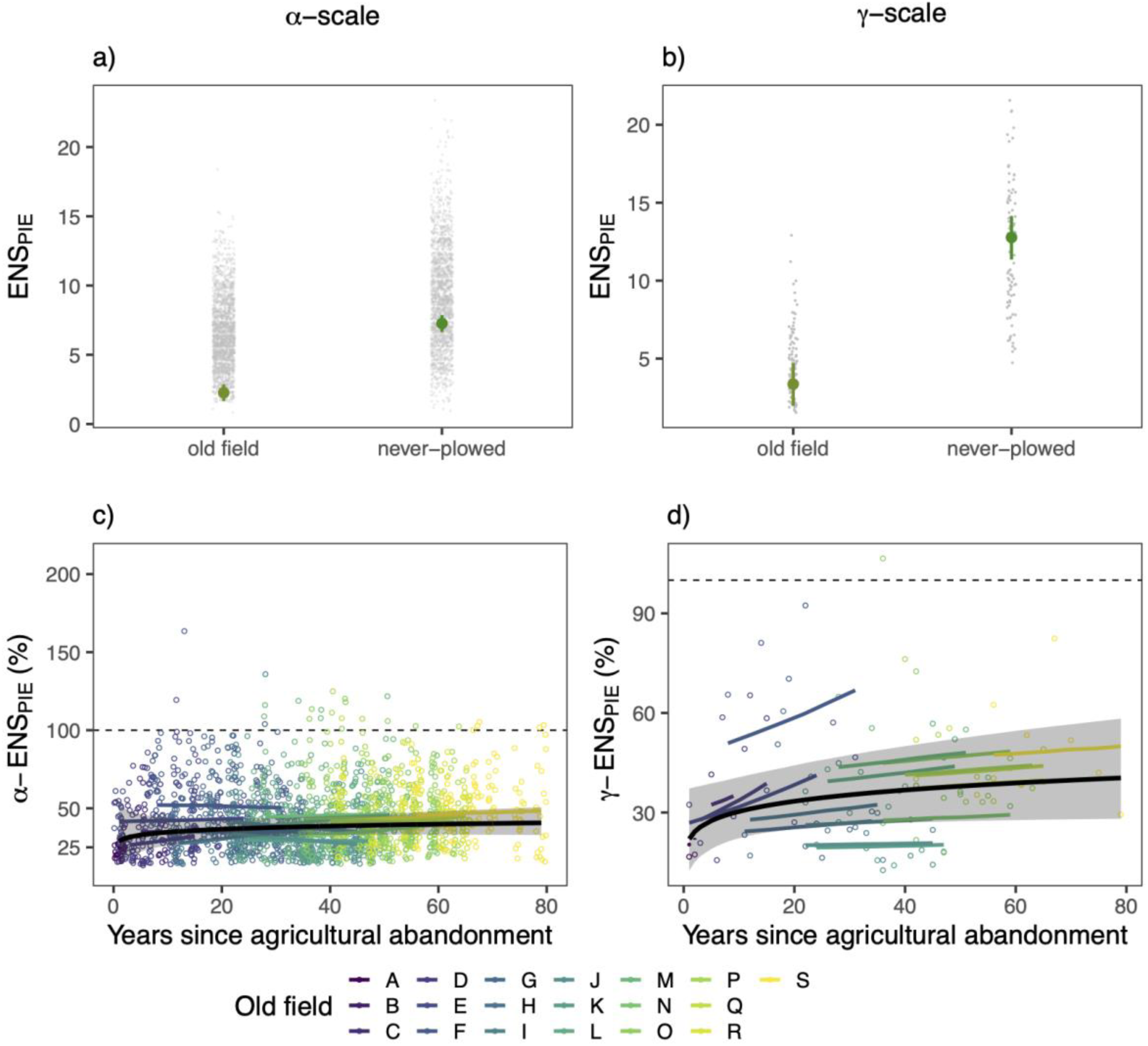
α-scale and γ-scale ENS_PIE_. a) α-scale and b) γ-scale ENS_PIE_ as a function of site status. Small points show data models were fit to; large points are the conditional effects of site status and the lines show the 95% credible intervals of conditional effects. c) α-scale and d) γ-scale species evenness as a function of ‘years since agricultural abandonment’. Black dashed line represents the mean diversity metric of all never-ploughed sites (18 sites). The thick black line represents the mean fitted line of years since agricultural abandonment on species richness across all old-fields. The grey shading around the black line represents the 95% credible interval of that mean effect estimate. Each colored line represents the average predicted values for each site. Each open point represents an old-field plot (α-scale) or a site (γ-scale) calculated as the percentage of evenness, compared to never-ploughed sites. Y-axes vary for clarity.

### β-diversity

In the discrete analysis, β-diversity (β = γ/α, Whittaker, 1972) values were 82% (3.50, 95% Credible Interval: 3.25 to 3.77) of that found in never-ploughed sites (4.28, 4.02 to 4.53, Fig. 3a). In the continuous analysis, ß-diversity values increased notably across years since agricultural abandonment (Slope: 0.15, CI: 0.09 to 0.22), and recovered up to 90% of the heterogeneity of that compared to never-ploughed sites (Fig. 3b).

In the discrete analysis, β-ENS_PIE_ values were 82% (1.31, CI: 1.15 to 1.46) of that found in never-ploughed sites (1.59, CI: 1.45 to 1.75, Fig. 3c). In the continuous analysis, β-ENS_PIE_ did not increase across years since agricultural abandonment (Slope: 0.10, CI: 0.02 to 0.19, Fig. 3d), and recovered to about 95% of that compared to never-ploughed sites.

**Figure 3:**
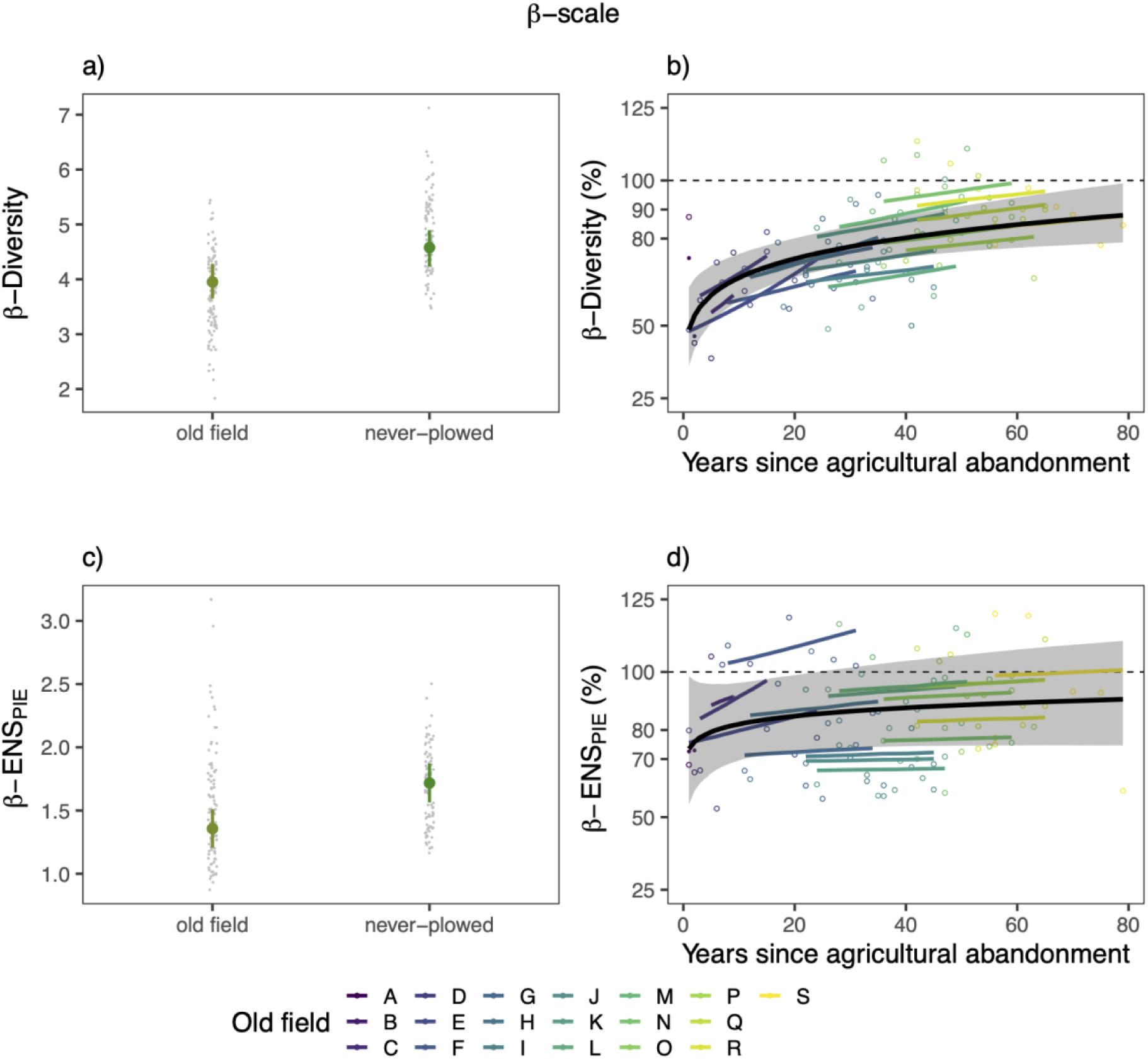
Whittaker’s β-Diversity (a, b) and β-ENS_PIE_ (c, d). a) & c) as a function of site status and b) & d) as a function of ‘years since agricultural abandonment’. In a) and c) small points show data models were fit to; large points are the conditional effects of site status and the lines show the 95% credible intervals of conditional effects. In b) and d) the thin black dashed line represents the mean diversity metric of all never-ploughed sites (18 sites). The thick black line represents the fitted mean line of years since agricultural abandonment on each diversity metric across all old-fields. The grey shading around the black line represents the 95% credible interval of that mean effect estimate. Each colored line represents the average predicted values for each site. Each open point represents an old-field calculated as the percentage of β-diversity, or β-ENS_PIE_ compared to the overall average of never-ploughed sites. Each colored line shows the slope of each site across years since agricultural abandonment. Y-axes vary for clarity.

### Community Composition

Dissimilarity due to turnover in old-fields compared to the never-ploughed region decreased across years since abandonment (Slope: -0.02, 95% Credible Interval: -0.03 to -0.01) (Fig. 4). That is, old-fields were colonised with species that were unique to old-field sites, when first abandoned, and this decreased over time. Dissimilarity due to nestedness in old-fields compared to never-ploughed sites increased across years since abandonment (Slope: 0.01, CI: 0.001 to 0.02) (Fig. 4). In other words, old-fields were increasingly colonised by species characteristic of the never-ploughed sites over time since abandonment, but never fully converged. In total, we found 63 species that occurred only in never-ploughed sites, never in any old-fields. Conversely, we found 34 species that occur only in old fields, and never in never-ploughed sites.

**Figure 4:**
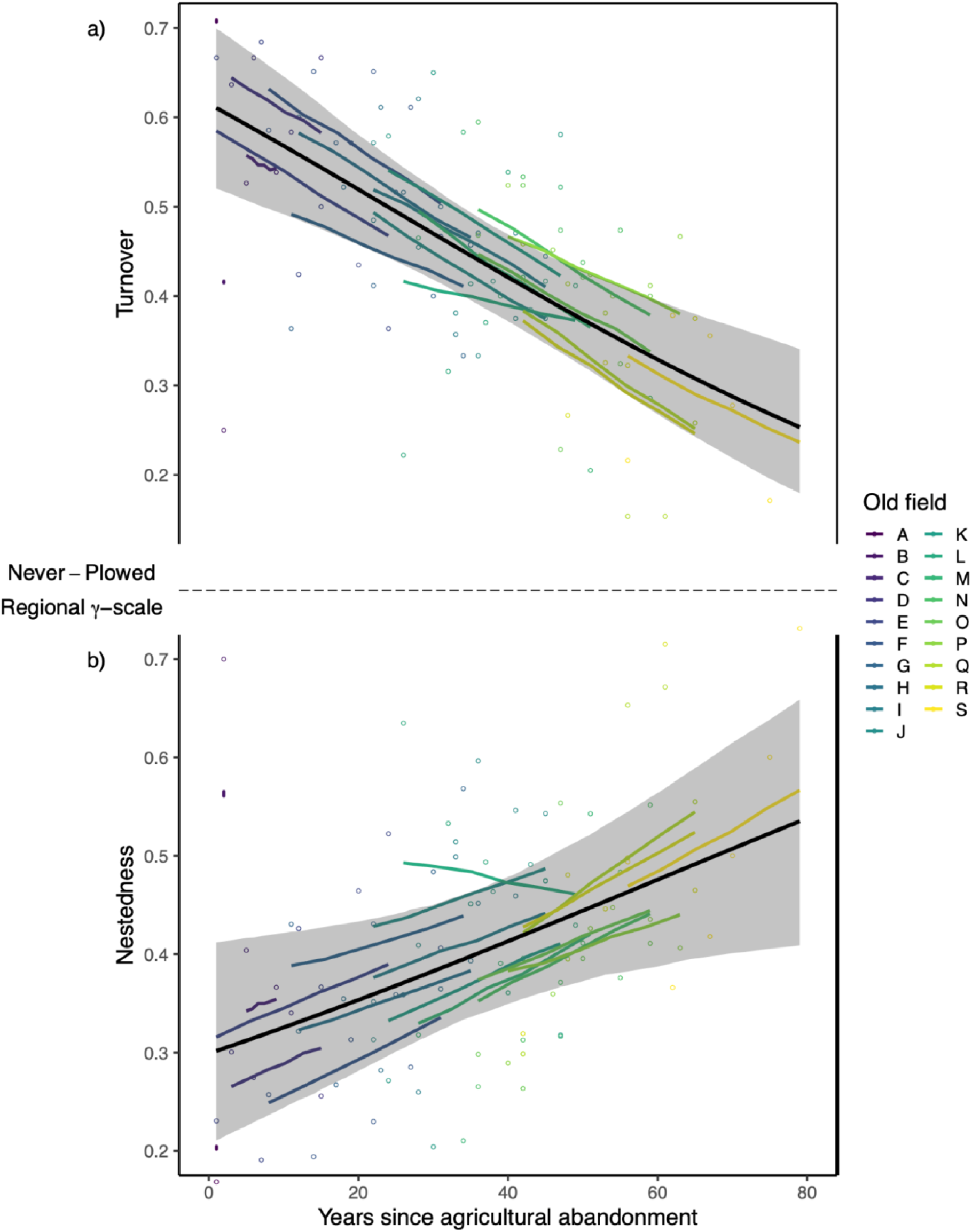
a) Spatial turnover and b) spatial nestedness components of Jaccard’s dissimilarity index as a function of ‘years since agricultural abandonment’. Each old-field at each time point was compared to the regional-γ species pool of all never-ploughed sites. The black dashed line represents a value of zero (0). The thick black line represents the average effect of years since agricultural abandonment on components across all old-fields compared to never-ploughed sites. The grey shading around the black line represents the 95% credible interval of that effect estimate. Each open point represents an old-field at a time since agricultural abandonment compared to the never-ploughed region. Each colored line shows the predicted slope of each old-field across years since agricultural abandonment.

### Growth Form and Origin Cover

Relative cover of growth form groups (graminoid, forb, legume) of differing origins (i.e., native and introduced) found in never-ploughed fields changed as a function of years since agricultural abandonment in old fields (Fig. 5). Native graminoid cover increased during years since agricultural abandonment in many sites, but the credible intervals overlap zero (Slope: 0.15, CI: -0.50 to 0.85). Both native forbs (-0.70, CI: -1.15 to -0.27) and native legumes (-1.34, CI: -2.36 to -0.36) decreased. Introduced graminoid cover increased in many sites since abandonment but again the credible intervals overlap zero (0.29, CI: -0.12 to 0.67). Both introduced fobs decreased (-0.57, CI: -0.98 to -0.20) and introduced legumes decreased (-0.10, CI: -0.86 to 0.56).

**Figure 5:**
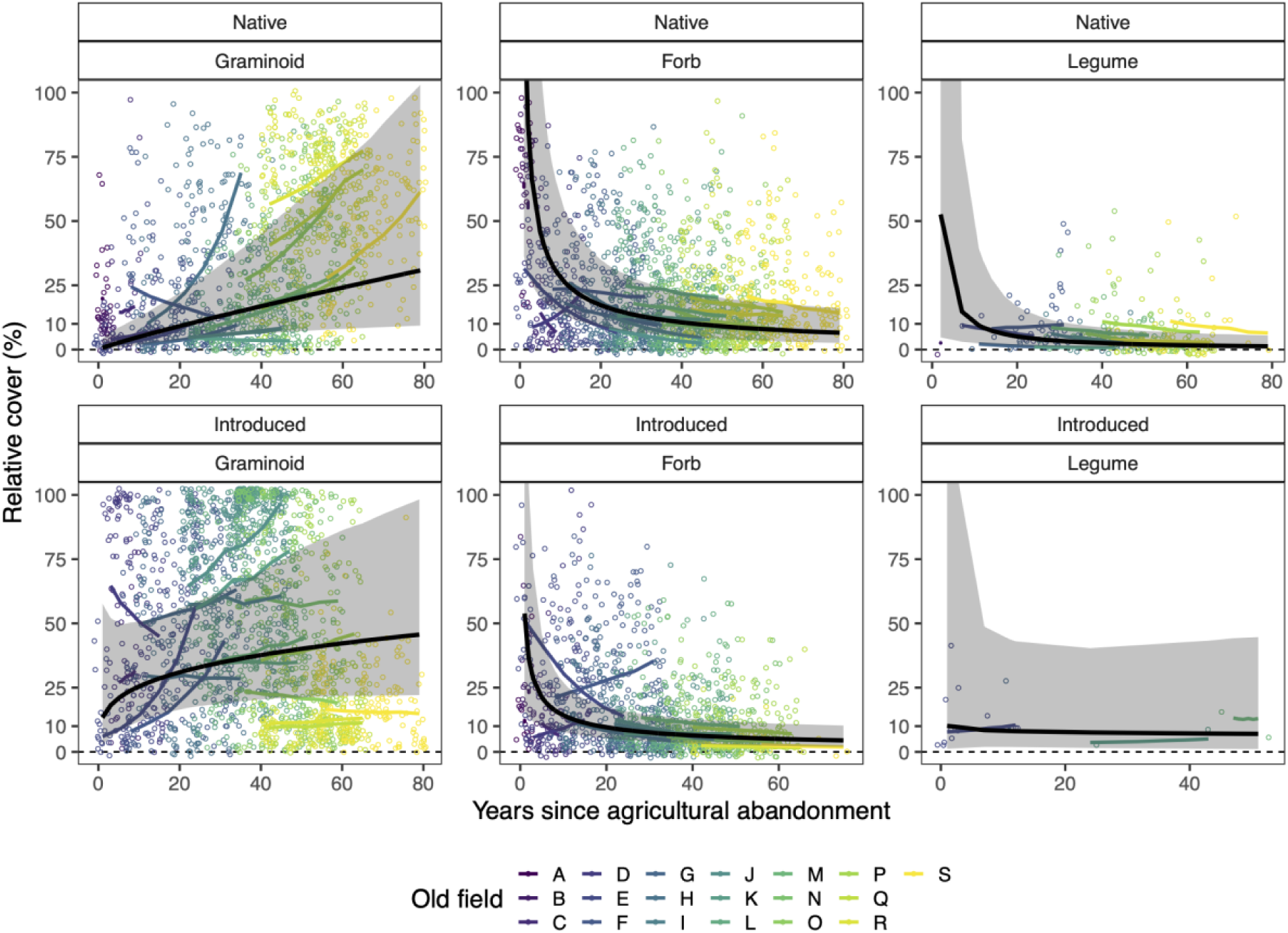
Relative cover of native a) graminoids b) forbs and c) legumes, and introduced d) graminoids e) forbs and f) legumes, characteristic of never-ploughed sites as a function of ‘years since agricultural abandonment’. The thick black line represents the average effect of years since agricultural abandonment on species growth form and origin groups across all old-fields. The grey shading around the black line represents the 95% credible interval of that effect estimate. Each open point represents the relative cover of each respective growth form and origin in an old-field plot at a time since agricultural abandonment. Each colored line shows the predicted slope of each old-field across years since agricultural abandonment for each growth form and origin.

## Discussion

A major tenet in disturbance ecology has been that, given enough time, removal of major anthropogenic disturbances such as agriculture will allow biodiversity to recover (Moreno-Mateos et al., 2017). At the same time, accruing evidence suggests that, in the absence of active restoration interventions (and even in their presence), this recovery can take an exceedingly long time and is often incomplete (Buisson et al., 2018; Isbell et al., 2013; Nerlekar & Veldman, 2020). For example, using many of the same sites as in the present study (see Supplementary Table 1 for details), Isbell et al. (2019) showed that α-scale (0.3m^2^) species richness had recovered to only ∼75% of that of the never-ploughed sites even after 91 years of recovery. Not surprisingly, we found similar results in our α-scale (0.5 m^2^) analyses, where within-plot species richness increased slightly through time, but remained ∼70% lower than never-ploughed plots, even after 80 years of recovery (Fig. 1c). While passive recovery is a nice option given resource constraints, it carries with it some hidden costs, such as incomplete recovery (Zahawi et al., 2014). Here, we examine the scale-dependent dynamics of this passive recovery over long-time scales. Given that active restoration is more expensive upfront, is not silver-bullet solution (Bekessy et al., 2010), nor is yet predictable (Brudvig & Catano, 2021), perhaps understanding scale-dependent dynamics of passive recovery better can help point to actionable improvements for restoration.

Importantly, coexistence and diversity are highly scale-dependent patterns and it is less clear how larger-scale patterns of diversity recover. Anthropogenic disturbances are known to often influence β-diversity in grassland ecosystems, both positively and negatively (Catano et al., 2017; Eskelinen & Harrison, 2015; Martin et al., 2005; Polley et al., 2005). As a result, we would either expect exacerbated effects of that driver and recovery at larger spatial scales (if β-diversity is lower) or enhanced effects of that driver and recovery at larger spatial scales (if β-diversity is higher). We found that β-diversity was indeed lower than in never-ploughed sites following agricultural abandonment, and while this diversity showed signs of recovery over time, it remained lower overall than in the never-ploughed sites. Complementary results have been found in successional sites that showed heterogeneity among plots increasing, and across sites decreasing, as succession progressed (S. Li et al., 2016). Other studies found that fire treatments increased in α-diversity, while β-diversity remained unchanged (Joner et al., 2021).

Given the lower β-diversity in recovering compared to never-ploughed sites (Fig 3a), we might expect that the magnitude of the deficit of species richness in recovered relative to never-ploughed sites might increase with spatial scale. This was true for the absolute number of species; the deficit in recovering relative to never-ploughed sites was ∼4 species at the α-scale (Fig. 1a), but ∼19 species at the γ-scale (Fig. 1b). But it was not true for the ratio of the deficit; the deficit of species richness was 59% at the α scale (Fig. 1a), with a similar recovery deficit of 55% at the γ-scale (Fig. 1b). In addition, there was little evidence that the deficit in species richness declined through time over the course of the observations, consistent with comparable findings and speculations on the slow recovery of secondary grasslands (Nerlekar & Veldman, 2020).

Although we cannot explicitly discern the mechanism underlying the continued deficit of species richness at α and γ-scales in these recovering old-fields, we suspect dispersal limitation between sites might play a key role. Several previous studies at Cedar Creek have shown that seed additions can lead to significantly increased levels of species richness (Fargione et al., 2011; Ladouceur et al., 2020; Symstad, 2000; Tilman, 1997). Even seeds added to a never-ploughed prairie-savanna led to a doubling of α plant diversity that persisted for 13 years or longer (Catford et al., 2019; Tilman, 1997). For example, the γ-diversity of Field H recovers rapidly (Fig. 1d), and is located right next to field N which has been recovering after abandonment for 35 years more than field H (Fig. S1, Table S1). This is evidence for spatially-dependent dispersal limitation. This is also consistent with our results comparing the nestedness and turnover components of compositional dissimilarity, which indicated that species composition in the old-fields was becoming more similar to the never-ploughed sites over time (Fig. 4), likely through species gains (Foster & Tilman, 2000). It is also possible that the soils in recovering sites have been significantly altered by added fertilisers so that environmental filters also play a role in limiting recovery (Seabloom et al., 2020).

By comparing the results of how species richness recovered following agricultural abandonment to those of ENS_PIE_, a diversity metric that strongly weights the most common species, we can see how patterns of recovery are influenced by more common versus rare species. In this case, ENS_PIE_ following agricultural abandonment recovered to only ∼31% of that observed in the never-ploughed sites at the α-scale (Fig. 2a). Compared to the ∼59% recovery of species richness at the α-scale (Fig. 1a), this suggests that much of the recovery was among species that are relatively common in the community and that there is less recovery of community evenness. At the site scale, ENS_PIE_ in old-fields recovered to 28% of that in never disturbed sites (Fig. 2b), compared to the 55% recovery of species richness at the site scale (Fig. 1b). The higher recovery of species richness than ENS_PIE_ is also consistent with previous studies (Martin et al., 2005; Sluis, 2002; Wilsey et al., 2005), suggesting that rarer species drive a lot of the passive recovery in abandoned old-fields.

Despite the fact that measurements of species diversity (richness and evenness) only partially recovered across scales even after nearly a century of agricultural abandonment, species composition has consistently recovered through time (Fig. 4). Early in the time series, recovering old-fields were colonised by species not typically found in never-ploughed sites. Grass cover characteristic of never-ploughed fields, both native and exotic, increased in old fields through time, and forb cover declined, suggesting that some growth forms may predictably recolonise (Clark et al., 2019), and increase their cover more readily than others (Fig. 5). Similar studies in other systems that have observed that perhaps species that are the slowest to recover on their own are less-effective dispersers (Fensham et al., 2016). Finally, we predict that without intervention, recovery to 95% of reference sites may take much longer, but this time to recovery may be different for each scale (Fig. S10). Given that successional recovery has been found to be most successful in colder, more humid systems (Prach & Walker, 2019), passive recovery, and even active restoration in more arid and hot systems is expected to be more difficult (Shackelford, Paterno, et al., 2021).

Here, to actively assist recovery, the control of invasive grasses, combined with direct seeding of native forbs, and management actions (e.g., fire, soil restoration) may be a favourable action to accelerate the recovery of diversity. Additionally, focusing restoration treatments on specific native grass and forb species that have not recovered may help to prioritise resources and actions through time. Combined with appropriate management actions (Guiden et al., 2021), and targeted support for trophic relationships (Heelemann et al., 2012; Ladouceur et al., 2022), restoration can also have cascading effects for passively supporting the recovery of the fauna community structure and function (Pearson et al., 2022).

Overall, our results show that analyses of multiple metrics across scales more fully reveals how ecological communities recover following disturbance in space and time. To accelerate or assist this recovery, active intervention via restoration may be considered a viable option. Understanding how biodiversity recovers on its own after disturbance across space and time can help us to better assist this recovery and restore systems more effectively and predictably into the future.

## Acknowledgements

We thank the staff, researchers, students, and interns who have worked for more than three decades to maintain the old field surveys and experiments at Cedar Creek. Data collection was supported by the NSF LTER program, including DEB- 8114302, DEB- 8811884, DEB-9411972, DEB-0080382, DEB-0620652, and DEB-1234162, and by the Cedar Creek Ecosystem Science Reserve and the University of Minnesota. We gratefully acknowledge the support of the German Centre for Integrative Biodiversity Research (iDiv) Halle-Jena-Leipzig, funded by the German Research Foundation (DFG–FZT 118, 202548816). We thank Christian Krause and the UFZ administrative and support staff of the High-Performance Computing Cluster EVE, a joint effort of the Helmholtz Centre for Environmental Research (UFZ) and iDiv, for access to, and support associated with, EVE. EL thanks the Alexander von Humboldt Foundation. Authors declare no conflict of interest.

## Author Contributions

EL, JMC and FI conceived the idea. FI and ATC contributed ideas and suggestions for data selection, preparation, analysis and interpretation. GDT, PBR and ATC provided data. EL conducted data analysis. JMC & WSH contributed ideas to analyses. EL & JMC wrote the manuscript. EL, FI, ATC, WSH, PBR, GDT and JMC contributed to the shaping of the manuscript, and made edits and suggestions leading to the final version.

## Data Availability

Data is archived at Cedar Creek Ecosystem Science LTER Reserve Data Catalogue: https://www.cedarcreek.umn.edu/research/data. Data summaries at different scales needed to reproduce results will be archived at a data repository like DRYAD or FigShare. Code will be archived at https://github.com/emma-ladouceur and assigned a DOI through Zenodo.

