## Supplementary Information for "The recovery of plant community composition following passive restoration across spatial scales"

***The recovery of biodiversity and species composition after agricultural abandonment across spatial scales***

**Table S1:** Site details. Range of years sampled, range of years since agricultural abandonment (YSA), number of years in which sampling occurred (# Years), mean species richness for each site at each scale (α-scale, ß-scale, γ-scale) (Cedar Creek E133 & E014). All fields used here were also used in (Isbell et al., 2019) but were part of a different set of surveys. Site 600 and 601 used in Isbell et al. 2019 were not used here as they were not included in surveys used in these studies. For more information on each experiment, see: https://www.cedarcreek.umn.edu/research/data

| Study | Site | Range of Years | Range of YSA | # Years | α-scale | ß-scale | γ-scale |
|---|---|---|---|---|---|---|---|
| 14 | A (601) | 2016-2016 | 1-1 | 1 | 2.85 | 4.21 | 12 |
| 14 | B (600) | 2016-2016 | 2-2 | 1 | 3.3 | 2.12 | 7 |
| 14 | C (10) | 2002-2006 | 5-9 | 4 | 6.38 | 2.74 | 16.5 |
| 14 | D (28) | 1994-2006 | 3-15 | 12 | 5.03 | 3.31 | 16.5 |
| 14 | E (41) | 1983-2006 | 1-24 | 23 | 7.36 | 3.1 | 22.17 |
| 14 | F (39) | 1983-2006 | 8-31 | 23 | 8.82 | 3.04 | 26.67 |
| 14 | G (40) | 1983-2006 | 11-34 | 23 | 6.51 | 3.51 | 22.83 |
| 14 | H (4) | 1983-2006 | 12-35 | 23 | 6.12 | 3.64 | 22.17 |
| 14 | I (44) | 1983-2006 | 22-45 | 23 | 5.67 | 3.19 | 18 |
| 14 | J (53) | 1983-2006 | 22-45 | 23 | 4.36 | 3.47 | 15 |
| 14 | K (47) | 1983-2006 | 24-47 | 23 | 5.63 | 4.15 | 23.17 |
| 14 | L (21) | 1983-2006 | 26-49 | 23 | 6.26 | 3.18 | 19.83 |
| 14 | M (70) | 1983-2006 | 28-51 | 23 | 7.79 | 4.39 | 33.67 |
| 14 | N (5) | 1983-2006 | 36-59 | 23 | 5.45 | 4.77 | 25.83 |
| 14 | O (27) | 1983-2006 | 36-59 | 23 | 8.19 | 3.92 | 31.83 |
| 14 | P (45) | 1983-2006 | 40-63 | 23 | 6.97 | 3.75 | 25.83 |

| | | | | | | | |
|---|---|---|---|---|---|---|---|
| 14 | Q (32) | 1983-2006 | 42-65 | 23 | 6.42 | 4.35 | 28.17 |
| 14 | R (35) | 1983-2006 | 42-65 | 23 | 6.43 | 4.66 | 30 |
| 14 | S (72) | 1983-2006 | 56-79 | 23 | 7.89 | 4.15 | 32.5 |
| 133 | 1 | 1984-2010 | never-plowed | 26 | 11.18 | 4.53 | 50.83 |
| 133 | 11 | 1984-2010 | never-plowed | 26 | 5.8 | 6.25 | 36.4 |
| 133 | 13 | 1984-2010 | never-plowed | 26 | 5.66 | 4.53 | 25.4 |
| 133 | 16 | 1984-2010 | never-plowed | 26 | 9.42 | 5.37 | 50.4 |
| 133 | 17 | 1995-2010 | never-plowed | 15 | 7.34 | 4.94 | 36.75 |
| 133 | 19 | 1990-2010 | never-plowed | 20 | 8.86 | 5.18 | 45.8 |
| 133 | 24 | 1995-2010 | never-plowed | 15 | 9.45 | 4.04 | 38 |
| 133 | 3 | 1984-2010 | never-plowed | 26 | 10.5 | 3.95 | 41.33 |
| 133 | 4 | 1984-2010 | never-plowed | 26 | 8.89 | 4.79 | 42.33 |
| 133 | 5 | 1984-2010 | never-plowed | 26 | 11.52 | 3.83 | 43.67 |
| 133 | 6 | 1990-2010 | never-plowed | 20 | 10.21 | 4.73 | 48 |
| 133 | 7 | 1984-2010 | never-plowed | 26 | 9.67 | 5.09 | 49 |
| 133 | 8 | 1984-2010 | never-plowed | 26 | 7.49 | 5.3 | 39.8 |
| 133 | 901 | 1995-2010 | never-plowed | 15 | 10.75 | 4.6 | 49.5 |
| 133 | 902 | 1995-2010 | never-plowed | 15 | 6.86 | 5.49 | 37.5 |
| 133 | 903 | 1990-2010 | never-plowed | 20 | 11.25 | 4.47 | 49.8 |
| 133 | 904 | 1995-2010 | never-plowed | 15 | 9.06 | 5.2 | 47.25 |
| 133 | 905 | 1990-2010 | never-plowed | 20 | 9.01 | 4.8 | 43.2 |

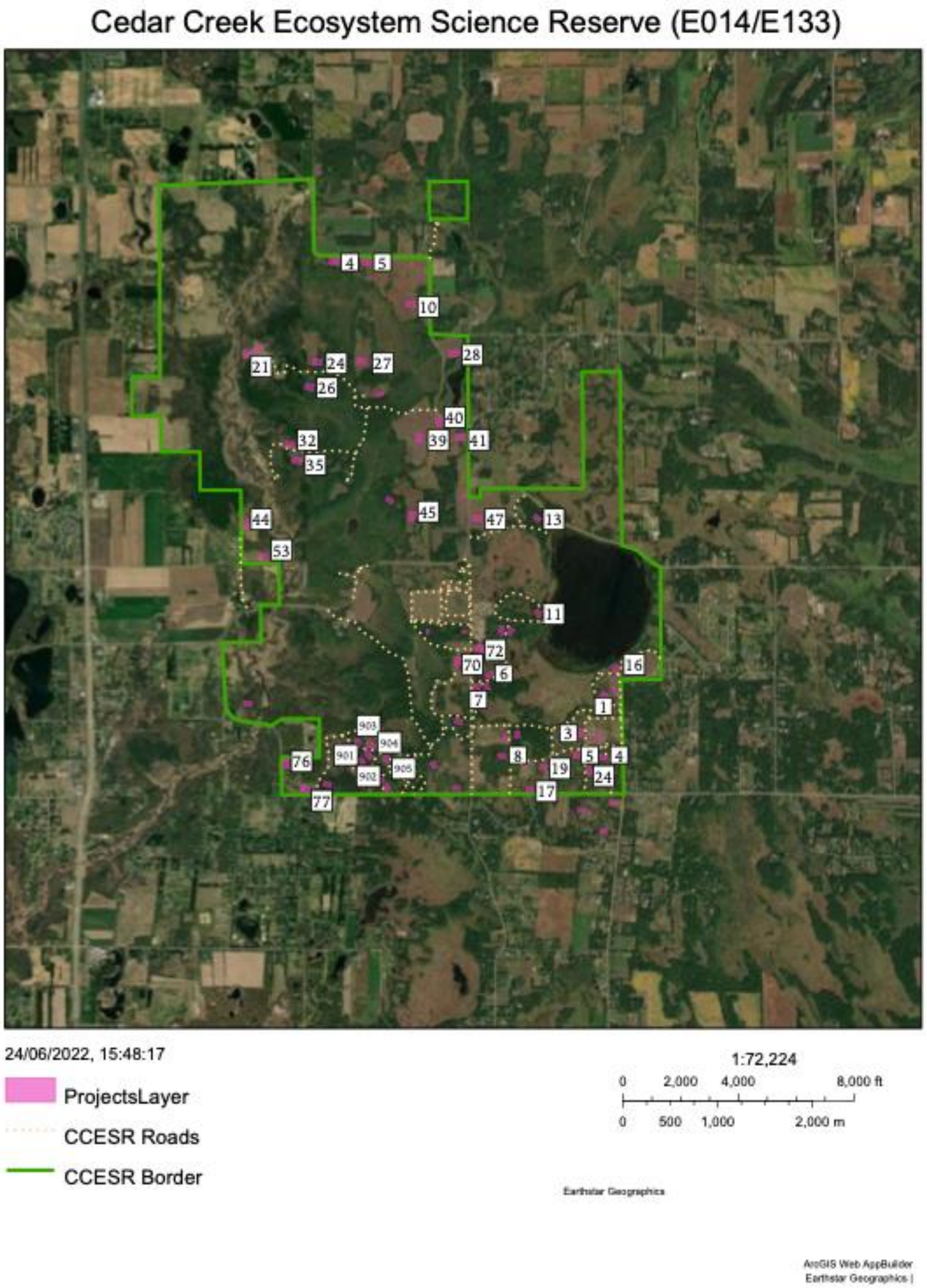


**Figure S1:** A map of sites, indicated by pink polygons, from Experiment 014 & 133 at Cedar Creek Ecosystem Science Reserve. Sites labelled with numbers were included in this study.

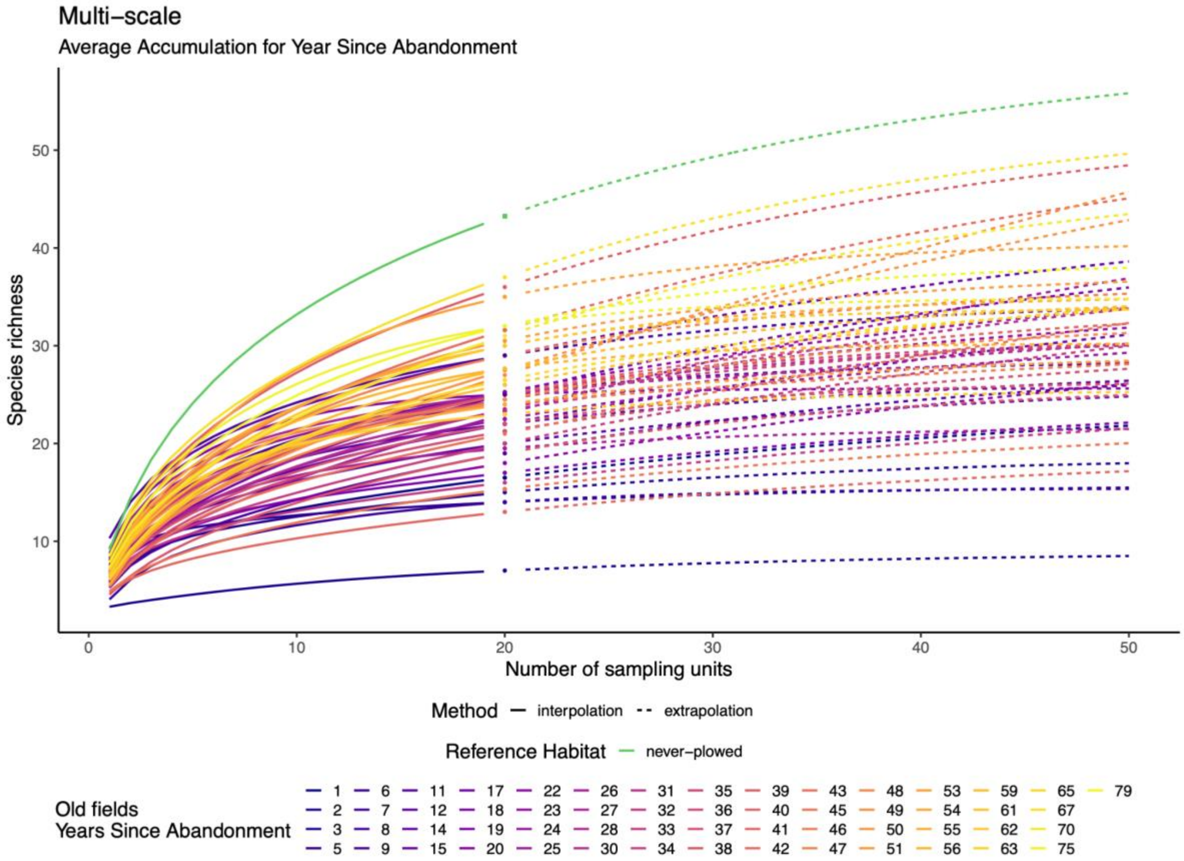


**Figure S2:** Incidence-based species accumulation curves across 20 sampling units as an average for each Year since abandonment, and an average across all never-ploughed sites. Solid lines indicate interpolated values, points indicate average observed values at 20 samples, and dotted lines indicate extrapolated values from 21 to 50 samples.

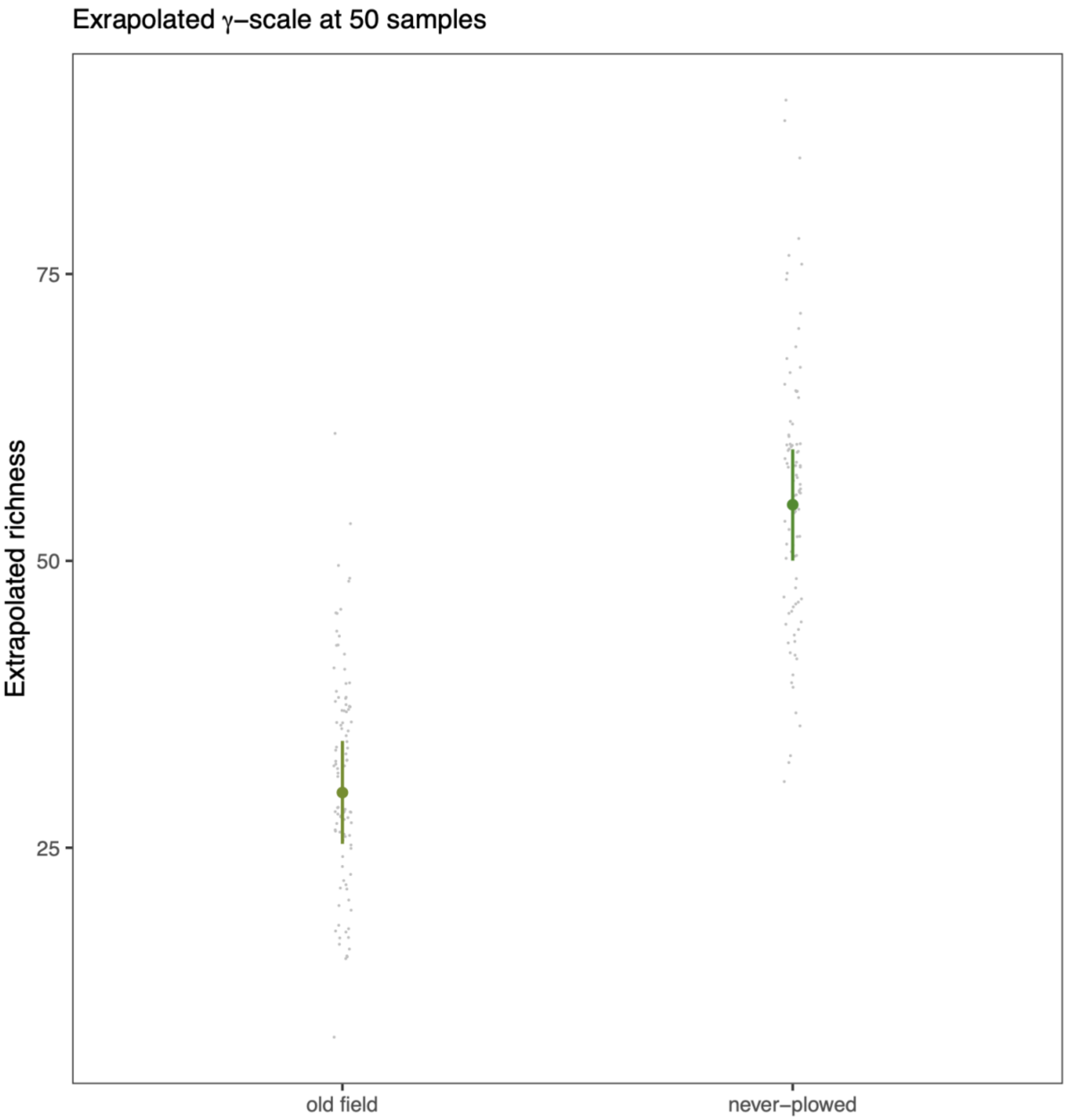


**Figure S3:** Extrapolated species richness estimated for 50 samples as a function of field status. Small points show data models were fit to; large points are the conditional effects of field status and the lines show the 95% credible intervals of conditional effects.

#### Supplementary Information

### Model Details

**Statistical Models**
**Discrete Models**
Responses were fit to ‘site-status’ (old-field, or never-ploughed) as a categorical fixed effect. Random intercepts and slopes were allowed to vary for each individual field and categorical calendar year.

**Continuous Models**
Responses were fit to ‘Year since agricultural abandonment’ as a continuous fixed effect (1-79 years). Random intercepts and slopes were allowed to vary for each individual field and categorical calendar year.

**Dissimilarity Models**
Turnover and nestedness components of Jaccard’s dissimilarity index were fit to ‘Year since agricultural abandonment’ as a continuous fixed effect. Random intercepts were allowed to vary for each individual field and categorical calendar year. Random slopes were allowed to vary for every field, and categorical calendar year.

**Table S2: Model specifications and performance**
1000 iterations were used as warm-up for all models. Model performance was confirmed by the visual inspection of predicted (light grey lines) values vs. observed (black line), and the distribution of residuals across all considered effects in each model for each response listed below. If a model reproduced the data well but had low sample sizes, we increased model sampling incrementally by 1000 iterations until large effective sample sizes, low Rhats and model convergence were achieved.

| Model | Figure | Distribution | Iterations | Effective Sample Size | Rhat |
|---|---|---|---|---|---|
| Discrete Models | | | | | |
| $\alpha$-scale species richness | 1 a) | Student | 3000 | 5810-1132 | ≤ 1.00 |
| $\gamma$-scale species richness | 1 b) | Poisson (log-link function) | 2000 | 1257-758 | ≤ 1.00 |
| $\alpha$-scale $ENS_{PIE}$ | 2 a) | Student | 3000 | 5608-1149 | ≤ 1.00 |
| $\gamma$-scale $ENS_{PIE}$ | 2 b) | Student | 3000 | 2850-1516 | ≤ 1.00 |
| $\beta$-scale Diversity | 3 a) | Student | 10000 | 27067-6662 | ≤ 1.00 |
| $\beta$-scale $ENS_{PIE}$ | 3 c) | Student | 4000 | 6666-2664 | ≤ 1.00 |

Continuous models

| | | | | | |
|---|---|---|---|---|---|
| $\alpha$-scale species richness (%) | 1 c) | Student | 10000 | 23002-1646 | ≤ 1.00 |
| $\gamma$-scale species richness (%) | 1 d) | Student | 3000 | 8617-229 | ≤ 1.00 |
| $\alpha$-scale $ENS_{PIE}$ (%) | 2 c) | Student | 7000 | 19886-763 | ≤ 1.01 |
| $\gamma$-scale $ENS_{PIE}$ (%) | 2 d) | Gaussian | 12000 | 29797-3812 | ≤ 1.00 |
| $\beta$-scale Diversity (%) | 3 b) | Student | 6000 | 13507-1819 | ≤ 1.00 |
| $\beta$-scale $ENS_{PIE}$ (%) | 3 d) | Student | 10000 | 20301-3351 | ≤ 1.00 |

Dissimilarity models

| | | | | | |
|---|---|---|---|---|---|
| Turnover | 4 a) | Zero-one-inflated-beta (logit-link function) | 2000 | 5338-453 | ≤ 1.00 |
| Nestedness | 4 b) | Zero-inflated-beta (logit-link function) | 2000 | 4108-541 | ≤ 1.00 |

Relative cover of growth forms

| | | | | | |
|---|---|---|---|---|---|
| Relative cover | 5 | Student | 2000 | 728-1716 | ≤ 1.00 |

Extrapolated species richness (50 samples)

| | | | | | |
|---|---|---|---|---|---|
| Extrapolated Richness | S3 | Student | 3000 | 5574-5093 | ≤ 1.00 |

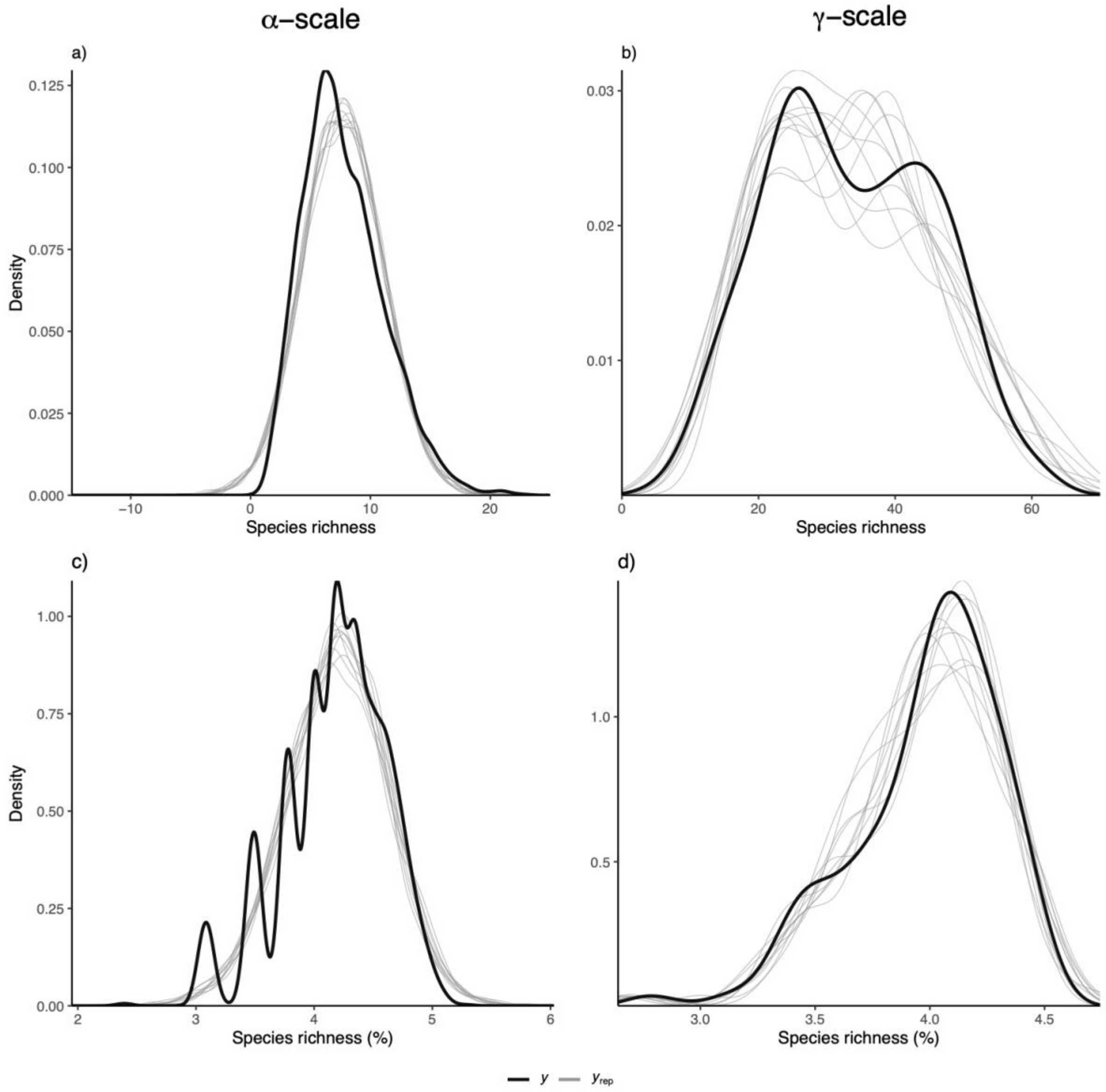


**Figure S4 a-d:** Predicted values for each model indicated by the grey lines and observed values of the data indicated by the black line for species richness for discrete models a) at the α-scale and b) at the γ-scale, and for continuous models c) at the α-scale and d) at the γ-scale. (Associated with Figure 1)

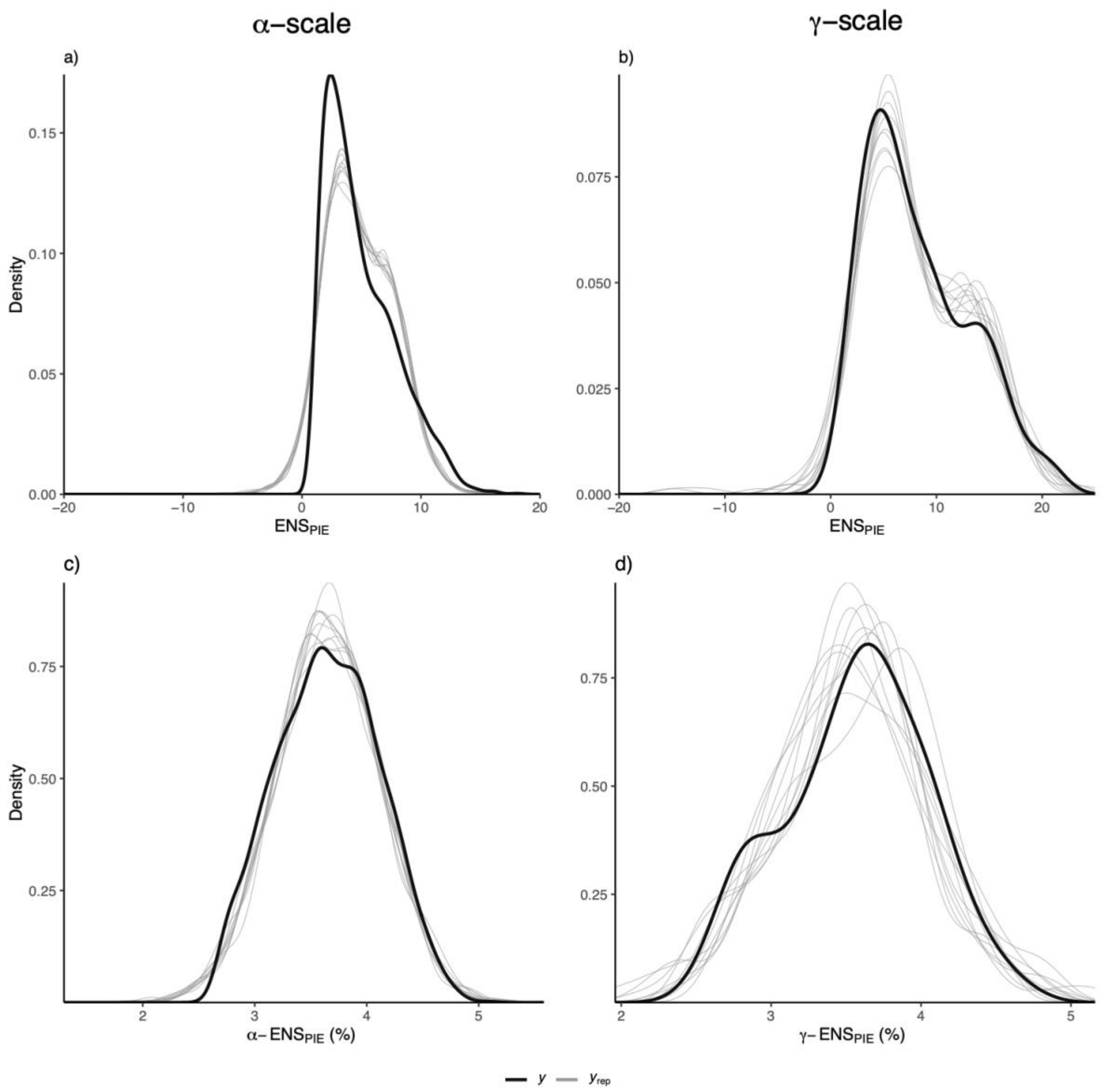


**Figure S5 a-d:** Predicted values for each model indicated by the grey lines and observed values of the data indicated by the black line for $ENS_{PIE}$ for discrete models a) at the $\alpha$-scale and b) at the $\gamma$-scale, and for continuous models c) at the $\alpha$-scale and d) at the $\gamma$-scale. (Associated with Figure 2)

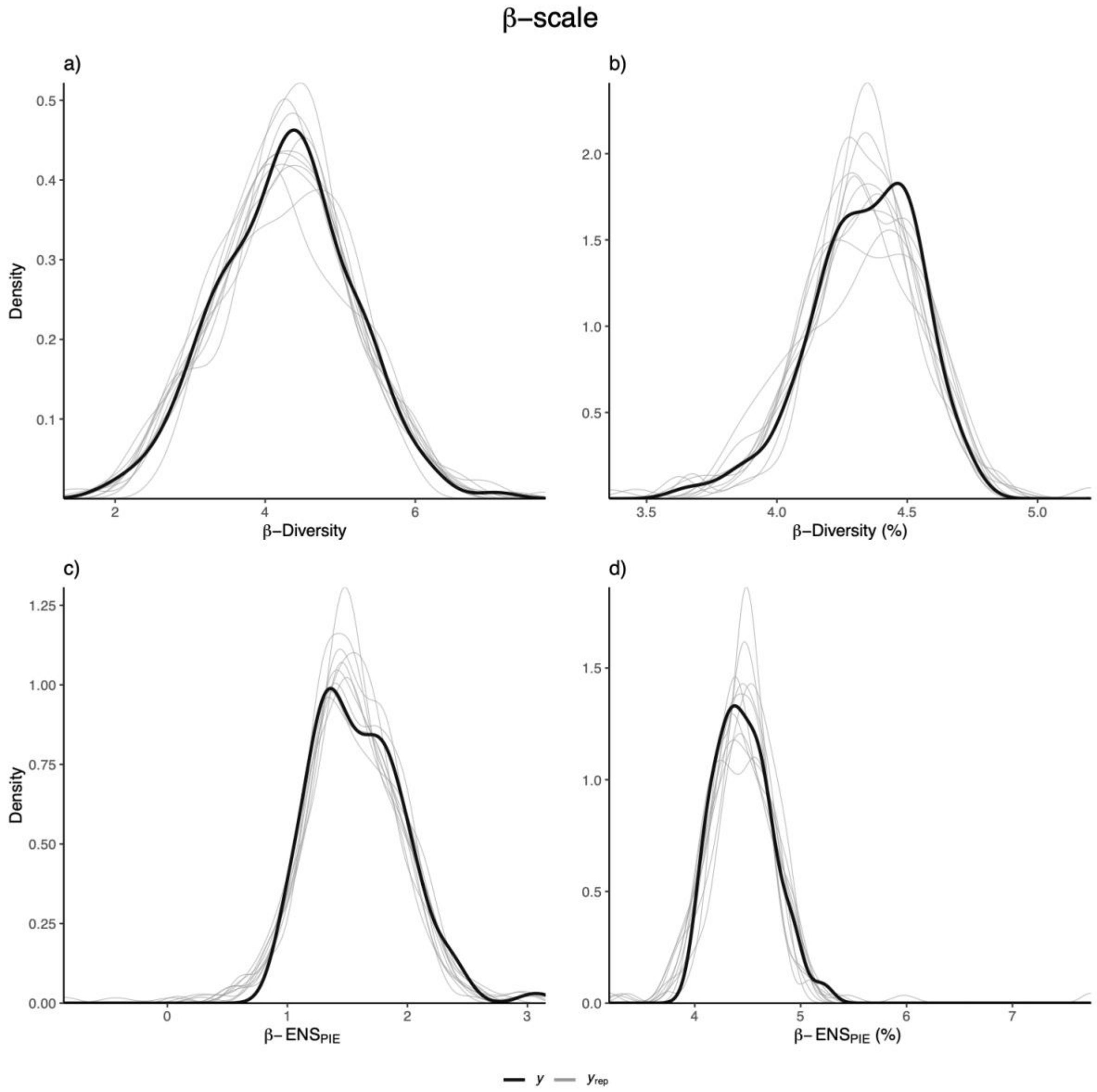


**Figure S6 a-d:** Predicted values for each model indicated by the grey lines and observed values of the data indicated by the black line at the β-scale for β-Diversity for a) discrete model and b) continuous model and for β- $ENS_{PIE}$ for the c) discrete model and b) continuous model. (Associated with Figure 3)

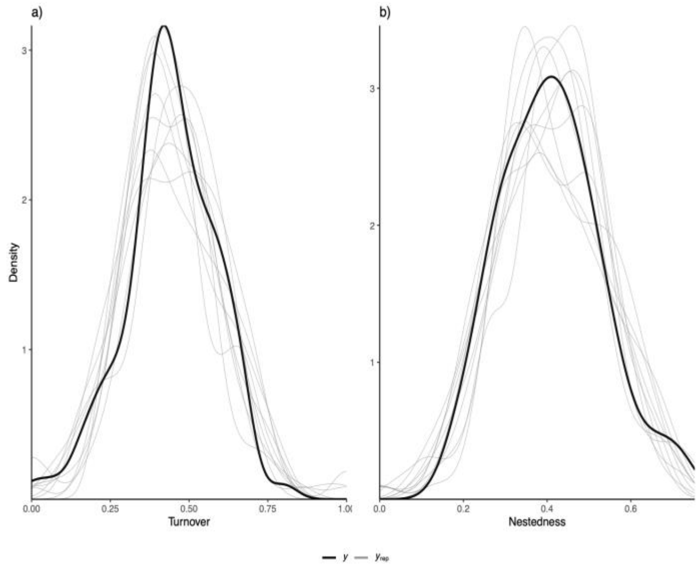


**Figure S7 a-b:** Predicted values for each model indicated by the grey lines and observed values of the data indicated by the black line of Jaccard's Dissimilarity Index for the a) Turnover component and b) the Nestedness component. (Associated with Figure 4)

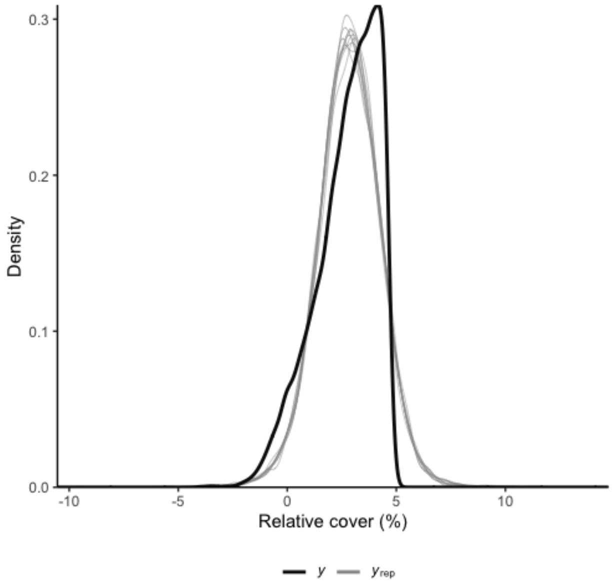


**Figure S8:** Predicted values for each model indicated by the grey lines and observed values of the data indicated by the black line of Relative cover of functional groups and their origin (Associated with Figure 5).

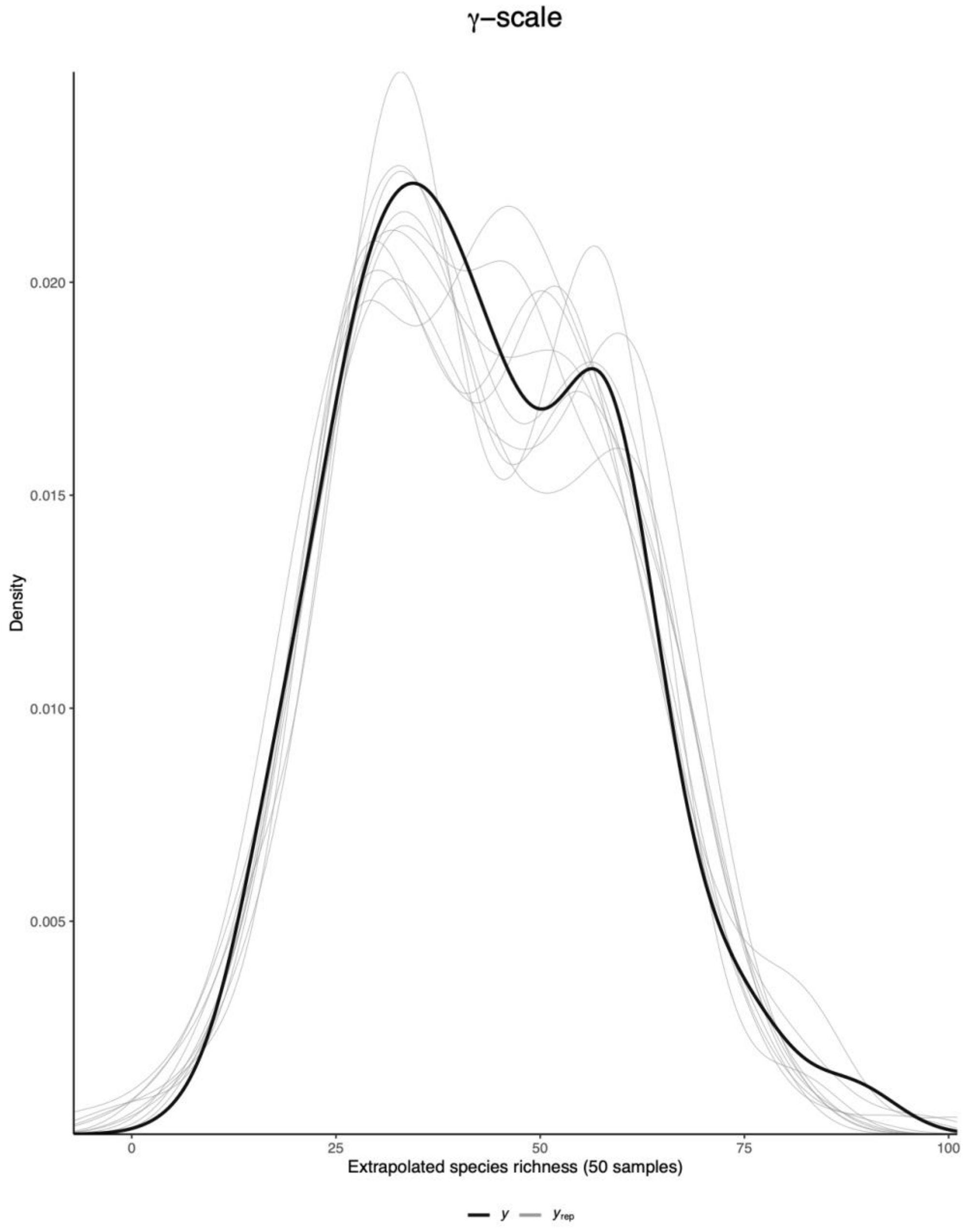


**Figure S9:** Predicted values indicated by the grey lines and observed values of the data indicated by the black line for extrapolated species richness for 50 samples at the γ-scale (Associated with Figure S3)

**Predicted Recovery**

Using the same models as for the continuous analyses for α, β, γ species richness (Figure 1c, 1d & 3b), we then predicted overall estimates for species richness across each scale (α-scale, γ-scale, β-scale) as function of year since agricultural abandonment. This quantified the predicted time to ~95%-100 recovery for each scale of measurement. We predicted values in intervals of 25 years for each metric up to a maximum of 1,000 years. β-diversity was predicted from the β-diversity model, but also by using predicted α and γ values to calculate and estimate predicted β-diversity, and estimates were extremely similar. These predictions harbour a large amount of uncertainty, and represent predictions of recovery only for these sites, under only this type of disturbance (agricultural).

By predicting percent recovery across each scale as a function of years since agricultural abandonment, we quantify how long old-fields might take to recover across time without intervention (active restoration) (Figure S10). We project that at the α-scale species richness is expected to recover to approximately 95% after 1000 years of abandonment, that γ-scale species richness is expected to recover to approximately 95% after 750 years of abandonment, and that β-Diversity is expected to recover to 95% of that seen in never-ploughed sites after approximately 150 years. We predict that heterogeneity and γ-scale species richness might recover faster than species richness at α-scales. These predicted values represent a great deal of uncertainty.

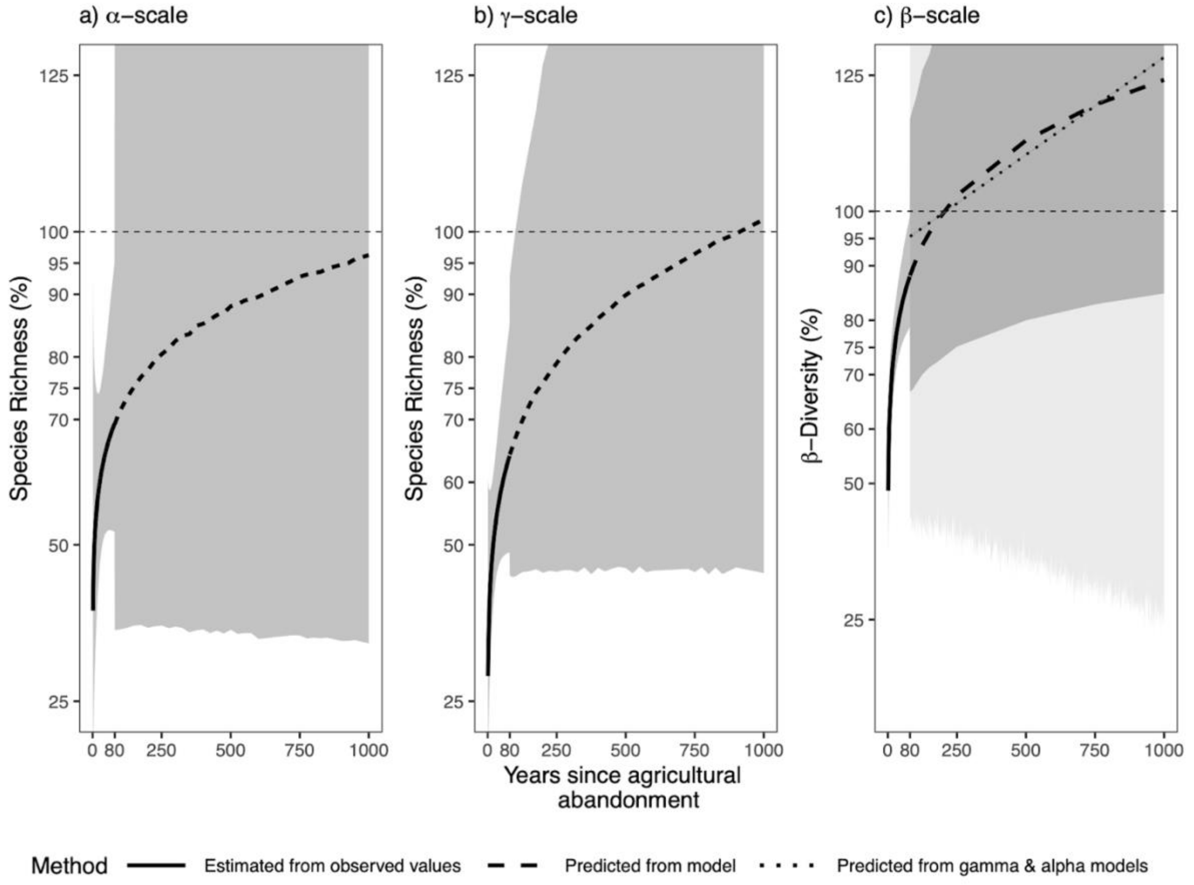


**Figure S10:** a) α-scale species richness b) γ-scale species richness c) and β-Diversity, % recovery as a function of 'years since agricultural abandonment'. Thin horizontal black dashed lines represent the mean diversity metric of all never-ploughed sites (18 sites). The thick black solid curved lines represent the average estimated effect of years since agricultural abandonment from observed values for each metric, and thick black dashed portions of the curve represent predicted estimates across years since abandonment. Dotted line in c) represents β-Diversity estimated from predicted α and γ values. The grey shading around the black line represents the 95% credible interval of that effect estimate for that model. The lighter grey shading in c) represents the uncertainty around the predicted estimate from gamma and alpha models.
